# Reduced Salience and Enhanced Central Executive Connectivity Following PTSD Treatment

**DOI:** 10.1101/501973

**Authors:** Chadi G. Abdallah, Christopher L. Averill, Amy E. Ramage, Lynnette A. Averill, Evelyn Alkin, John H. Krystal, John D. Roache, Patricia Resick, Stacey Young-McCaughan, Alan L. Peterson, Peter Fox, the STRONG STAR Consortium

## Abstract

**BACKGROUND:** In soldiers with posttraumatic stress disorder (PTSD), symptom provocation was found to induce increased connectivity within the salience network, as measured by functional magnetic resonance imaging (*f*MRI) and global brain connectivity with global signal regression (GBCr). However, it is unknown whether these GBCr disturbances would normalize following effective PTSD treatment.

**METHODS:** 69 US Army soldiers with (n = 42) and without PTSD (n = 27) completed *f*MRI at rest and during symptom provocation using subject-specific script imagery. Then, participants with PTSD received 6 weeks (12 sessions) of group cognitive processing therapy (CPT) or present-centered therapy (PCT). At week 8, all participants repeated the *f*MRI scans. The primary analysis used a region-of-interest approach to determine the effect of treatment on salience GBCr. A secondary analysis was conducted to explore the pattern of GBCr alterations posttreatment in PTSD participants compared to controls.

**RESULTS:** Over the treatment period, PCT significantly reduced salience GBCr (*p* = .02). Compared to controls, salience GBCr was high pretreatment (PCT, *p* = .01; CPT, *p* = .03) and normalized post-PCT (*p* = .53), but not post-CPT (*p* = .006). Whole-brain secondary analysis found high GBCr within the central executive network in PTSD participants compared to controls. *Post hoc* exploratory analyses showed significant increases in executive GBCr following CPT treatment (*p* = .01).

**CONCLUSION:** The results support previous models relating CPT to central executive network and enhanced cognitive control while unraveling a previously unknown neurobiological mechanism of PCT treatment, demonstrating treatment-specific reduction in salience connectivity during trauma recollection.

## Introduction

Advances in neuroimaging and connectomics have led to a shift in the clinical neuroscience field from an early focus on brain regions and localization to identifying neural circuits, and more recently, to establishing network functioning in health and disease (1). The investigation of neural correlates of posttraumatic stress disorder (PTSD) largely followed a comparable path. Early neuroimaging PTSD studies identified a number of regions of interest, which were then integrated into circuitry related hypotheses and more recently into network-based models (2–5). These network models suggested an association between PTSD and increased salience network but reduced default mode and central executive network connectivity (2,3,5). However, these models were primarily based on findings from seed analyses of resting-state functional connectivity magnetic resonance imaging (*fc*MRI) data in cross-sectional studies. Unfortunately, the seed-based approach does not fully interrogate the brain’s large-scale intrinsic connectivity networks (ICNs) (6). Additionally, the resting-state data may not necessarily generalize to functioning during provoked symptoms or other tasks (7). Moreover, the cross-sectional investigations are, by design, limited to association evidence without the ability to ascertain the network changes over the course of the illness. These limitations could be partially mitigated by employing graph-based measures and task *fc*MRI in longitudinal studies. Using a graph-based measure named global brain connectivity with global signal regression (GBCr), the current report complements previous literature by conducting a longitudinal *fc*MRI investigation at rest and during symptom provocation in active duty US Army soldiers with and without PTSD. The participants with PTSD were scanned pre-and post-randomized treatment with group cognitive processing therapy (CPT) or present-centered therapy (PCT).

Nodal strength (also known as nodal degree) is the amount of connections between a node and the nodes of the rest of the network. It is a fundamental measure in a graph-based network, as the majority of other network topology measures are ultimately related to it (8). Over the past decade, GBCr, a well-established measure of nodal strength, provided robust and reproducible evidence of network disturbances in several psychiatric disorders (9–17). GBCr was also found to be sensitive to treatment, with accumulating evidence of normalization of GBCr disturbances following ketamine treatment of depressed patients (9–11). In combat-exposed US military veterans, prefrontal GBCr did not correlate with PTSD total symptom severity (18). However, clusters of high prefrontal GBCr were found in those who reported high arousal over the past month (18). This raises the question whether the level of symptoms during the scan may have increased the GBCr values in this subpopulation. Recently, this hypothesis was supported by a data-driven cross-sectional analysis demonstrating increased GBCr within the salience network during symptom provocation, but not at rest, in PTSD compared to trauma and non-trauma control (19).

In this report, we investigated the longitudinal effects of psychotherapy on the GBCr alterations in the salience network during symptom provocation. This was accomplished by conducting a region of interest (ROI) analysis examining the effects of CPT and PCT on GBCr compared to a nontreated combat control (CC) group without PTSD. Then, using a previously established approach (9), we conducted a data-driven whole-brain analysis comparing post-treatment GBCr, during symptom provocation, between the PTSD group and CC. The aim of this approach is to identify patterns of normalization (i.e., absence of pretreatment disturbances) and adaptation (i.e., evidence of new alterations). Follow-up ROI analyses examined whether the posttreatment alterations were differentially influenced by CPT and PCT. The study predictions were that psychotherapy will significantly reduce salience GBCr, leading to a normalization pattern posttreatment.

## Methods

The behavioral and imaging data were provided by the STRONG STAR data repository (https://tango.uthscsa.edu/strongstar/subs/rpinfo.asp?prj=12). The clinical trial results for the PTSD treatment study were previously reported (20). The pre-treatment GBCr data were reported elsewhere (19). The posttreatment data and analyses are new and have not been reported previously.

### Study Population

PTSD (n = 42) and CC (n = 27) active military participants with successful scans were investigated (Table 1) as a subset of a larger randomized controlled clinical trial (20). The patients with PTSD completed pretreatment scans and received CPT (cognitive only (21)) or PCT (22) group therapy (90-minute sessions, twice per week for 6 weeks). Post-treatment scans were repeated 2 weeks after the end of treatment (i.e., a total of approximately 8 weeks between scans). Similarly, the CC group completed repeated scans, 8 weeks apart, without receiving any intervention.

**Table 1.** Demographics and Clinical Characteristics.

|  | PCT (n = 23) | CPT (n = 19) | Combat Control (n = 27) |
| --- | --- | --- | --- |
|  | Mean (SEM) or N (%) | Mean (SEM) or N (%) | Mean (SEM) or N (%) |
| Age | 32.7 (1.9) | 32.3 (1.5) | 31.8 (1.1) |
| BMI | 29.6 (1.0) | 28.4 (1.0) | 28.2 (0.6) |
| IQ | 97 (1.8) | 102 (2.6) | 99 (2.1) |
| Sex (Male) | 22 (96%) | 17 (90%) | 25 (93%) |
| Race (White) | 16 (70%) | 13 (68%) | 17 (63%) |
| Race (Black) | 4 (17%) | 3 (16%) | 6 (22%) |
| Ethnicity (Hispanic) | 6 (26%) | 3 (16%) | 9 (33%) |
| Baseline BDI-II | 28.0 (2.7) | 28.4 (2.5) |  |
| Delta BDI-II | 7.0 (1.9) | 7.3 (2.8) |  |
| Baseline BAI | 27.0 (3.0) | 24.0 (2.5) |  |
| Delta BAI | 8.3 (2.0) | 8.6 (2.1) |  |
| Baseline PCL | 56.7 (2.3) | 56.4 (2.5) |  |
| Delta PCL | 11.7 (2.7) | 9.7 (3.8) |  |
| Baseline PSS-I | 27.2 (1.5) | 28.3 (1.7) |  |
| Delta PSS-I | 6.1 (2.6) | 6.2 (2.7) |  |

No significant differences between subgroups. Abbreviations: BAI, Beck Anxiety Inventory; BDI-II, Beck Depression Inventory – II; BMI, body mass index; CPT, cognitive processing therapy; IQ, intelligence quotient; PCL, PTSD Checklist; PCT, present-centered therapy; PSS-I, PTSD Symptom Scale – Interview Version; SEM, standard error of means.

All participants completed informed consent prior to participation. Study procedures were approved by institutional review boards. All participants had no MR contraindication and had a negative drug screen on the day of the scan. The clinical trial criteria were previously reported (20). Briefly, patients with PTSD were active duty US Army soldiers, following deployment to or near Iraq or Afghanistan, who were 18 years or older with *DSM-IV* PTSD diagnoses and were stable on or off psychotropic medications for at least 6 weeks; they did not have imminent suicide or homicide risk, psychosis, or more than mild traumatic brain injury. The CC group endorsed a Criterion A traumatic event during deployment but did not have current PTSD. Severity of symptoms pretreatment and posttreatment (week 8) were assessed using the PTSD Symptom Scale – Interview Version (PSS-I) (23), PTSD Checklist (PCL) for *DSM-IV* (24), the Beck Depression Inventory-II (BDI-II) (25), and the Beck Anxiety Inventory (BAI) (26).

### FcMRI Acquisition & Processing

The acquisition parameters were previously reported (19). Briefly, each *f*MRI scan (voxel size = 2 x 2 x 3 mm; TR = 3000 ms; TE = 30 ms) included 10 minutes at rest and 12 minutes during symptom provocation – i.e., script imagery during which participants listened to recorded retelling of a personal event (alternating between trauma and neutral) over a 1-minute period, followed by a 1-minute period of thinking about the event and then a 1-minute break. The Human Connectome Pipeline was adapted to conduct surface-based preprocessing and optimize registration (27). Details of our image processing pipeline were previously reported (11) and are provided in the Supplemental Information. Following our previous reports (9–11,18), GBCr values were computed as the average of the correlations between each vertex/voxel and all other vertices/voxels in the brain gray matter (see Supplemental Information).

### Statistical Analyses

We used the Statistical Package for the Social Sciences (SPSS, version 24) for the behavioral and ROI analyses. The normal distribution of outcome measures was confirmed using probability plots and test statistics. The standard error of means (SEM) were provided as estimates of variation. Significance was set at *p* ≤ .05, with 2-tailed tests. ANOVA and chi square were used to compare behavioral data across groups.

To investigate the salience ROI (Fig. 1A; based on (19)), we constructed a general linear model (GLM) to determine the main effects of group (PCT vs. CPT vs. CC), task (rest vs. scripts), and time (pre-treatment vs. posttreatment), as well as the interactions between the main effects, followed by post-hoc pairwise comparisons.

**Figure 1.**
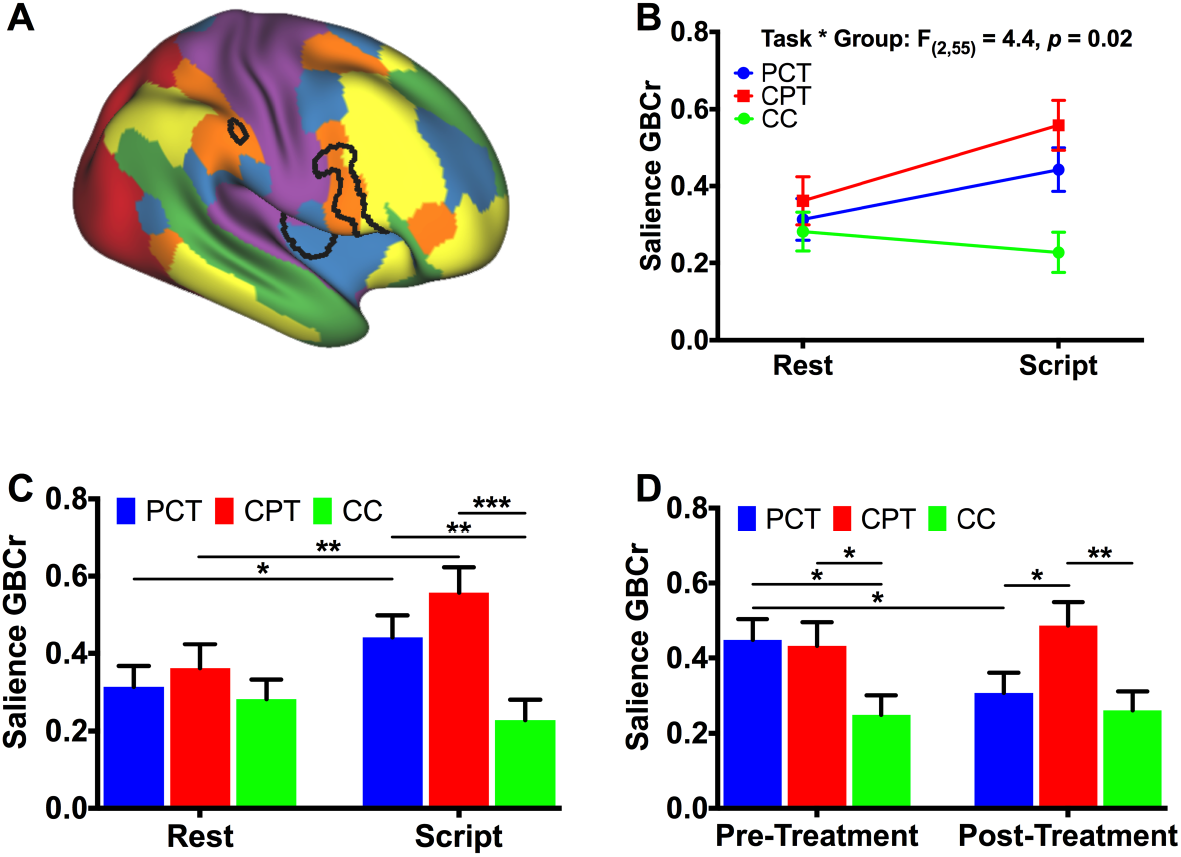
The Effects of Psychotherapy on Salience Connectivity. **A.** The map of 6 intrinsic connectivity networks: ventral salience (blue), dorsal salience (orange), central executive (yellow), default mode (green), visual (red), and sensorimotor (purple). The black lines mark the salience clusters based on previous cross-sectional findings. **B.** There was a significant group by task interaction effects on salience global brain connectivity with global signal regression (GBCr). **C.** There was significant increase in GBCr during trauma recollection (i.e., script imagery) compared to during resting state in posttraumatic stress disorder (PTSD) patients treated with present-centered therapy (PCT) or cognitive processing therapy (CPT), but not in combat control (CC). The higher GBCr values in PTSD compared to CC were significant only during trauma recollection, but not a rest. D. PCT, but not CPT, significantly reduced salience GBCr. * *p* ≤ .05; ** *p* ≤ .01; *** *p* ≤ .001.

To determine the pattern of GBCr alterations following treatment, we conducted a vertex-/voxel-wise *fc*MRI non-parametric analysis using FSL Permutation Analysis of Linear Models (PALM), with tail approximation and cluster mass threshold of 1.96 for Type I error correction (corrected *α* = .05) (28). This data-driven whole-brain analysis used independent *t* tests to identify posttreatment clusters with altered GBCr during symptom provocation in the PTSD group compared to CC. To facilitate the interpretation of the whole-brain findings (i.e., increase in executive GBCr), the identified clusters (vertex/voxel *p* < .005; corrected *α* = .05) were extracted to conduct follow-up post-hoc ROI analyses to better characterize the executive GBCr changes across time, tasks and subgroups. This was accomplished by conducting a GLM comparable to the one used for investigating the salience ROI.

Finally, we conducted exploratory analyses examining the correlation in the PTSD group between salience/executive GBCr and improvement/severity measures (BDI-II, BAI, PCL, PSS).

## Results

Participants were well matched for age, sex, body mass index (BMI), intelligence quotient (IQ), race, and ethnicity (Table 1). Pretreatment PSS-I, PCL, BDI-II, and BAI did not differ between treatment groups. In the clinical trial participants, both CPT and PCT significantly reduced clinical symptoms on the PSS-I, PCL, BDI-II, and BAI at week 8 (all *p* values < .05), but there were no significant differences between treatments (all *p* values > .6).

### Normalization: PCT Reduced Salience Functional Connectivity

Investigating the salience ROI (Fig. 1A), the GLM revealed significant effects of group (*F*(_2,55_) = 4.8, *p* = .01), task (*F*_(1,55)_ = 6.1, *p* = 0.02) and task*group interaction (*F*_(2,55)_ = 4.4, *p* = .02; Fig. 1B), with increased salience GBCr during symptom provocation compared to resting state in the PCT and CPT groups but not in CC (Fig. 1C). Additionally, salience GBCr values were higher in the PCT and CPT groups compared to CC during symptom provocation but not at rest (Fig. 1C). We also found trends for time*task (*F*_(1,55)_ = 3.4, *p* = .07) and time*group interaction (*F*_(2,55)_ = 2.8, *p* = .07; Fig. 1D), with significant reduction of salience GBCr following PCT (*p* = .02). Compared to CC, salience GBCr was high pretreatment for the 2 PTSD groups (PCT, *p* = .01; CPT, *p* = .03) and normalized post-PCT treatment (*p* = .53) but not post-CPT treatment (*p* = .006). There were no main time effects (*p* = .48) or time*task*group interaction (*p* = .75).

### Adaptation: CPT Enhanced Central Executive Functional Connectivity

In the participants with PTSD compared to the CC group, posttreatment whole-brain analysis revealed a significantly high GBCr in areas within the left ventrolateral prefrontal, right rostral-ventrolateral and dorsolateral prefrontal cortices (Fig. 2A-B). We also found significant clusters of low GBCr in the rostral-ventral areas of the cerebellum in the treated participants compared to the CC group (Fig. S1). Notably, the salience cluster, which showed high GBCr in the 2 treated groups compared to controls in the cross-sectional study (19), appears to normalize following treatment with adaptation shift toward higher GBCr within the central executive network (Fig. 2C-D). Hence, to facilitate the interpretation of the whole-brain findings, we conducted post hoc analyses by extracting average GBCr from each subject within this executive ROI, which included areas that showed significantly high GBCr in PTSD (Fig. 2).

**Figure 2.**
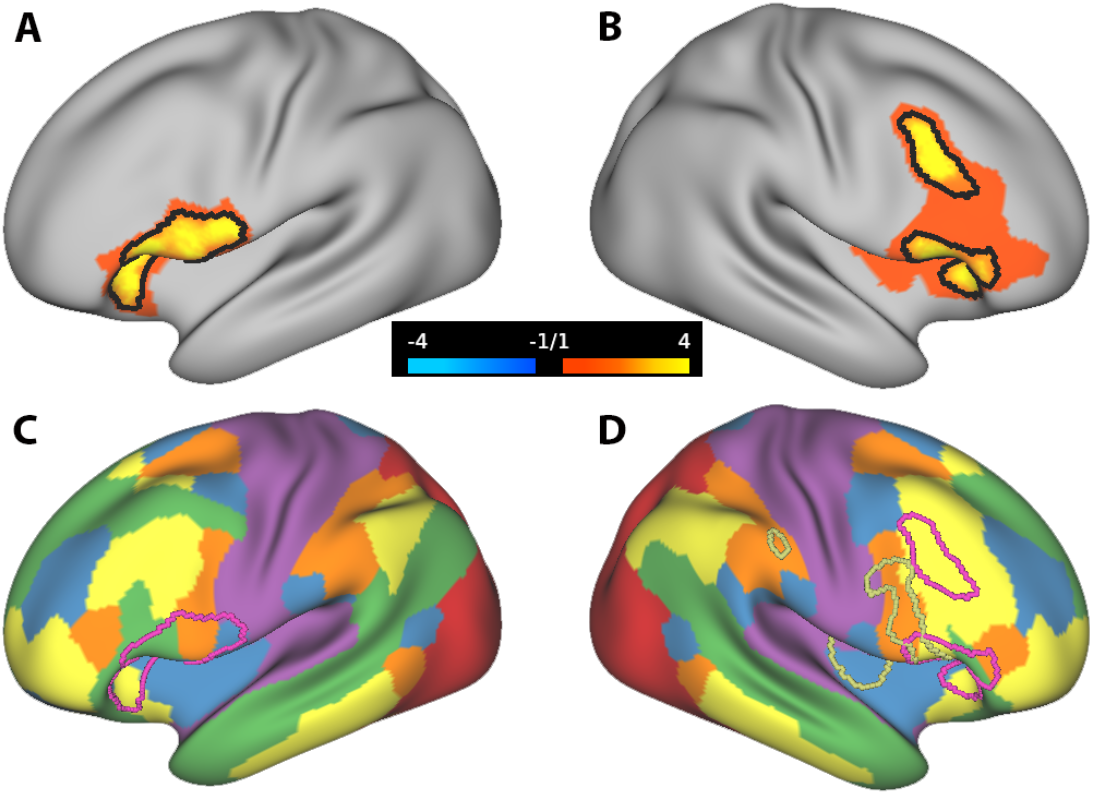
Cortical Global Connectivity Post-treatment. **A & B.** The red-yellow clusters mark the vertices with increased global brain connectivity with global signal regression (GBCr) in treated posttraumatic stress disorder (PTSD) compared to controls during symptom provocation. The black lines mark the vertices with *p* < .005 and corrected *α* = .05. **C & D**. The map of 6 intrinsic connectivity networks: ventral salience (blue), dorsal salience (orange), central executive (yellow), default mode (green), visual (red), and sensorimotor (purple). The dark-yellow lines mark the salience cluster and the red lines mirror the black lines in A & B, marking the executive cluster.

Investigating the executive ROI, the GLM revealed significant effects of group (*F*_(2,55)_ = 4.0, *p* =.02), task (*F*_(1,55)_ = 6.6, *p* = .01), and task*group (*F*_(2,55)_ = 7.1, *p* = .002; Fig. 3A-B) and task*time interaction (*F*_(1,55)_ = 7.3, *p* = .009; Fig. 3C), with increased executive GBCr during symptom provocation compared to resting state in the CPT and PCT groups but not in CC (Fig. 1C). Additionally, executive GBCr values were higher in the PCT and CPT groups compared to CC during symptom provocation but not at rest (Fig. 3B). We also found significant increases of executive GBCr following CPT (*p* = .01) during symptom provocation (Fig. 3D). There were no main time effects (*p* = .54), time*group interaction (*p* = .30), or time*task*group interaction (*p* = .22). Additional analyses of the cerebellar ROI are provided in the Supplemental Information (Fig. S2).

**Figure 3.**
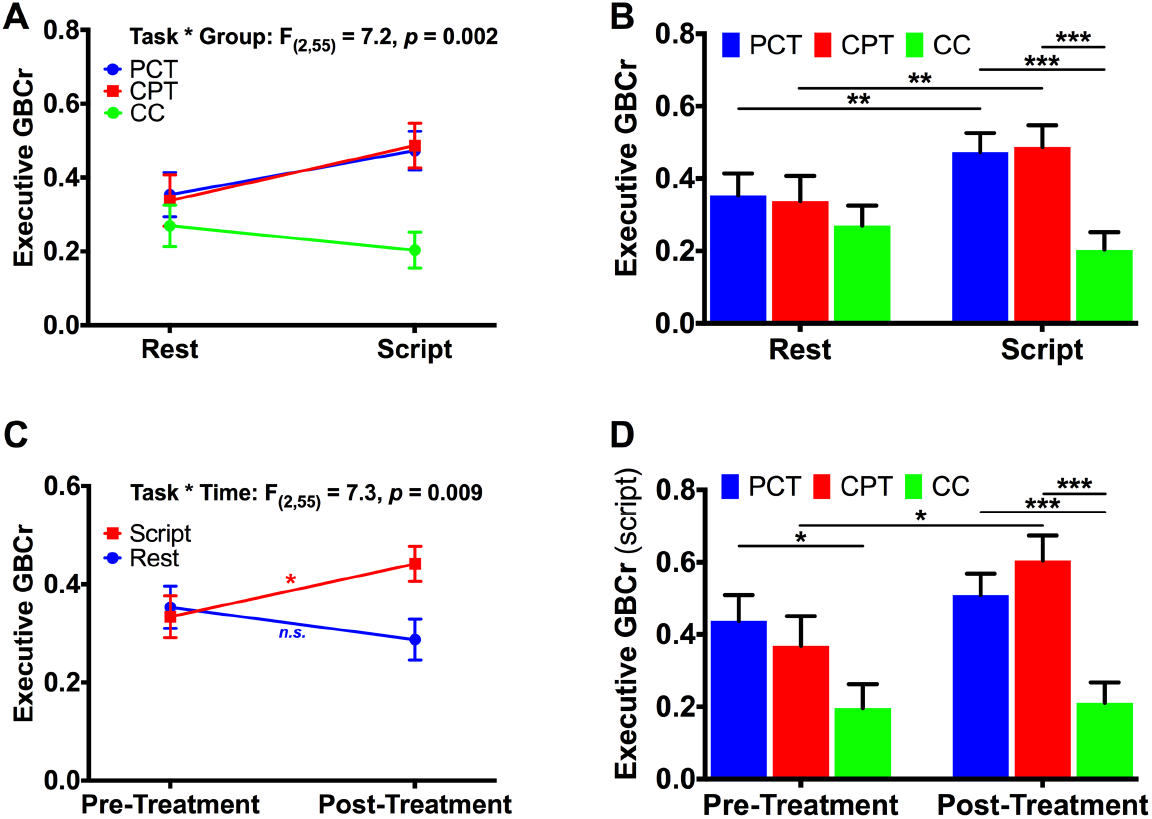
The Effects of Psychotherapy on Executive Connectivity. **A.** There was a significant group by task interaction effects on executive global brain connectivity with global signal regression (GBCr). **B**. There was significant increase in GBCr during trauma recollection (i.e., script imagery) compared to during resting state in post-traumatic stress disorder (PTSD) patients treated with present-centered therapy (PCT) or cognitive processing therapy (CPT), but not in combat control (CC). The higher GBCr values in PTSD compared to CC were significant only during trauma recollection, but not a rest. **C**. There was a significant time by task interaction effects on executive GBCr. **D**. CPT, but not PCT, significantly increased executive GBCr. *n.s*.: not significant; * *p* ≤ .05; ** *p* ≤ .01; *** *p* ≤ .001.

### Exploring the Relationship Between GBCr and Symptoms

Pretreatment executive GBCr during symptom provocation was associated with improvement in PCL scores (i.e., pre-minus post-treatment) over the treatment period (*r* = .36, *p* = .027). In addition, pre-treatment executive GBCr at rest was negatively associated with pretreatment BAI (*r* = –.42, *p* = .008), PCL (*r* = –.43, *p* = .006), and PSS-I scores (*r* = –.45, *p* = .004). No other correlations between executive GBCr and symptoms severity or improvement were found. We found no correlations between salience GBCr and improvement of symptoms. Finally, the readers should cautiously interpret these exploratory findings, considering that they do not survive correction for multiple comparisons.

## Discussion

Overall, we found a pattern of salience network normalization (i.e., reduction) and executive network adaptation (i.e., increase) following evidence-based psychotherapy in PTSD patients treated twice per week for 6 weeks. There were no significant cortical connectivity changes in the combat control group. Post hoc analyses showed that CPT induced a significant increase in executive connectivity leading to adaptation changes with higher salience and executive connectivity values post-CPT in PTSD compared to combat control. In contrast, PCT induced a significant reduction in salience connectivity leading to normalization and salience connectivity values comparable to combat control. Finally, the data-driven analysis posttreatment showed reduced global connectivity in PTSD in areas within the cerebellum, including both the spinocerebellum and cerebro-cerebellum (Fig. S1). However, there was no significant treatment effect compared to changes in combat control (Fig. S2).

CPT is a cognitive therapy in which patients examine their thinking and emotions about the traumatic event. The patients are systematically taught how to change their thinking to more balanced beliefs with the use of Socratic questioning by the therapist (20,21). The findings of CPT-related increases in global connectivity within the executive network are consistent with the cognitive model wherein executive control improves the processing of trauma-related stimuli, resulting in moderated expression of emotion in response to trauma-related cues. Consistent with this hypothesis, a previous study using seed-based analysis showed CPT-induced increases in central executive functional connectivity, which were interpreted as indicative of top-down cognitive control of affective processes that are disrupted in PTSD (29). Moreover, systemic reviews and meta-analyses of neuroimaging research have reported an association between cognitive therapies and increased activity in brain regions within the executive network (30,31). To further advance this hypothesis, future studies should investigate GBCr during a cognitive task to determine the extent of pretreatment executive abnormalities in PTSD and whether the connectivity of a cognitively engaged central executive network could predict response to psychotherapy or whether it is affected by treatment.

PCT was originally developed as an active comparator to trauma-focused cognitive therapy (22). Hence, PCT includes common components of efficacious psychotherapy without focusing on the trauma or using cognitive or supportive frameworks. PCT focuses on managing PTSD symptoms using psychoeducation and problem-solving strategies to generate possible solutions to current problems or PTSD symptoms that the patient can practice off-sessions. Although neuroimaging studies examining the effects of PCT are scarce, one pilot study reported PCT-induced reduction in the activation of the insula during the presentation of traumatic images and sounds (32). Another study reported increased resting state functional connectivity between clusters within the default mode network following PCT treatment (33). PTSD is associated with reduced default mode connectivity (6), an abnormality that is believed to be the result of an overactive salience network failing to effectively arbitrate between default mode and central executive networks (2,3). In this context, the previously reported PCT-induced reduction in insula activity during trauma cues and increase in default mode connectivity during resting state may reflect a pattern of normalization of the salience network following PCT treatment. The current study results further support this model by demonstrating significant reduction in trauma-induced global brain connectivity within the salience network following PCT. While the current study data do not allow us to distinguish which components of PCT are responsible for the reduction in salience connectivity, we speculate that perhaps the out-of-sessions, repeated practice of the symptom reduction solutions generated through problem-solving strategies during the sessions may have led to enhanced utilization of habitual rather than cognitive reactions to trauma cues.

Finally, accumulating evidence repeatedly demonstrates functional and structural abnormalities in the cerebellum of PTSD patients (34–39). Moreover, two recent studies have shown reduced functional nodal strength in the cerebellum in PTSD (19,40). In the current study, PTSD patients continued to show reduction in cerebellar connectivity posttreatment compared to controls (Fig. S1). Follow-up analyses showed persistently lower cerebellar connectivity in PTSD during trauma recollection, regardless of treatment modality, with no significant treatment effects compared to changes in combat control (Fig. S2).

### Limitations and Strengths

Considering that both interventions were active, efficacious treatments, the study design cannot confirm that the observed connectivity changes post-treatment are due to the specific intervention rather than generalized, nonspecific changes due to reduction in PTSD symptoms. However, the differential changes in connectivity patterns per treatment suggest a direct relationship between CPT and PCT with executive and salience global connectivity, respectively. Another limitation is that the executive ROI analyses are dependent on the vertex-wise results. Therefore, these data should be interpreted within the context of better understanding data-driven findings, rather than fully independent test results. Finally, the measure of nodal strength is not limited to a specific ICN, but rather measures the role of each node within the whole-brain network. Therefore, the salience and executive connectivity alternations may either indicate increased internal (i.e., within network) and/or external connectivity (i.e., between networks). Future studies could use network-restricted topology approaches (6) to further delineate the role of each ICN, as well as the interaction of ICNs.

The current study has many strengths including: (a) a longitudinal design in an adequate sample with randomization to 2 evidence-based efficacious treatments for these purposes; (b) the inclusion of repeated scans in the control group to account for nonspecific test-retest changes; (c) the use of symptom provocation paradigm to identify trauma-specific dynamic shift in ICNs; (d) the use of a well-validated measure of nodal strength. GBCr has been repeatedly associated with psychopathology and successful treatment (9–11). In addition, GBCr does not require *a priori* selection of seed or ROI, which here permitted the posttreatment data-driven analysis. In the current study, the lack of significant cortical GBCr changes in the control group underscores the robustness of the measure and the specificity of the study paradigm; (e) the use of state-of-the-art neuroimaging methods based on the Human Connectome Pipeline, including enhanced registration, surface-based analysis, and nonparametric correction for the vertex/voxel-wise multiple comparisons.

## Conclusions

The results provide strong neurobiological evidence supporting the role of the central executive network in the mechanism of CPT treatment to engage cognitive control and ultimately reduce PTSD symptoms. Intriguingly, the study findings may have unraveled a previously unknown neurobiological mechanism of PCT treatment, demonstrating treatment-specific reduction in salience connectivity during trauma recollection. It remains to be seen whether the normalized salience connectivity is primarily driven by the habitual reactions established through off-session practicing of symptom-reduction solutions devised during therapy sessions. In summary, evidence-based psychotherapy exerted a pattern of normalization within the salience network and adaptation in the executive network. While the adaptational changes favored CPT, the normalization was mostly limited to PCT. The used biomarkers are well-validated and have previously shown notable reproducibility following pharmacotherapeutic interventions (9–11). Therefore, future studies may capitalize on current findings to determine the clinical utility of these biomarkers in predicting or optimizing treatment for millions of patients suffering from PTSD.

## ACKNOWLEDGMENTS

The authors would like to thank the individuals who participated in these studies for their invaluable contribution. Funding for this work was made possible by grants to the STRONG STAR Consortium by the US Department of Defense through the US Army Medical Research and Materiel Command, Congressionally Directed Medical Research Programs, Psychological Health and Traumatic Brain Injury Research Program awards W81XWH-08-02-109 (Alan Peterson), W81XWH-08-02-0112 (Peter Fox), W81XWH-08-02-0114 (Brett Litz), and W81XWH-08-02-0116 (Patricia Resick). Some of the investigators also had additional support from the National Institute of Mental Health (K23MH101498) and the VA National Center for PTSD. The views expressed in this article are solely those of the authors and do not represent and endorsement by or the official policy or position of the Department of Defense, the Department of Veterans Affairs, the National Institutes of Health, or the US Government.

## CONFLICTS OF INTEREST

CGA has served as a consultant and/or on advisory boards for FSV7, Genentech and Janssen, and editor of *Chronic Stress* for Sage Publications, Inc.; he has filed a patent for using mTOR inhibitors to augment the effects of antidepressants (filed on August 20, 2018). JHK is a consultant for AbbVie, Inc., Amgen, Astellas Pharma Global Development, Inc., AstraZeneca Pharmaceuticals, Biomedisyn Corporation, Bristol-Myers Squibb, Eli Lilly and Company, Euthymics Bioscience, Inc., Neurovance, Inc., FORUM Pharmaceuticals, Janssen Research & Development, Lundbeck Research USA, Novartis Pharma AG, Otsuka America Pharmaceutical, Inc., Sage Therapeutics, Inc., Sunovion Pharmaceuticals, Inc., and Takeda Industries; is on the Scientific Advisory Board for Lohocla Research Corporation, Mnemosyne Pharmaceuticals, Inc., Naurex, Inc., and Pfizer; is a stockholder in Biohaven Pharmaceuticals; holds stock options in Mnemosyne Pharmaceuticals, Inc.; holds patents for Dopamine and Noradrenergic Reuptake Inhibitors in Treatment of Schizophrenia, US Patent No. 5,447,948 (issued September 5, 1995), and Glutamate Modulating Agents in the Treatment of Mental Disorders, U.S. Patent No. 8,778,979 (issued July 15, 2014); and filed a patent for Intranasal Administration of Ketamine to Treat Depression. U.S. Application No. 14/197,767 (filed on March 5, 2014); US application or Patent Cooperation Treaty international application No. 14/306,382 (filed on June 17, 2014). Filed a patent for using mTOR inhibitors to augment the effects of antidepressants (filed on August 20, 2018). All other co-authors declare no conflict of interest.

## SUPPLEMENTAL INFORMATION

### Image Processing

The Human Connectome Project (HCP) Pipelines (github.com/Washington-University/Pipelines) were adapted to process the imaging data (1). Briefly, the adapted minimal preprocessing included FreeSurfer automatic segmentation and parcellation of high-resolution structural scans, deletion of first 5 volumes, slice timing correction, motion correction, intensity normalization, brain masking, and registration of *f*MRI images to structural MRI and standard template, while minimizing smoothing from interpolation. Then, the cortical gray matter ribbon voxels and each subcortical parcel were projected to a standard Connectivity Informatics Technology Initiative (CIFTI) 2mm grayordinate space. ICA-FIX was run to identify and remove artifacts (2,3), followed by mean grayordinate time series regression (MGTR; which is comparable to global signal regression in volume data). The latter two processing steps (FIX+MGTR) have been found to significantly reduce motion-correlated artifacts (4). In addition, there were no differences (*p* > .1) in head motion during *f*MRI session between the study groups at rest (mean ±*SEM*; Pretreatment: PCT = .10 ± .018; CPT = .09 ± .007; CC = .07 ± .004; Posttreatment: PCT = .07 ± .008; CPT = .07 ± .005; CC = .06 ± .003) and during symptoms provocation (Pretreatment: PCT = .09 ± .006; CPT = .12 ± .019; CC = .09 ± .009; Posttreatment: PCT = .13 ± .031; CPT = .10 ± .007; CC = .10 ± .012).

Details of global brain connectivity with global signal regression (GBCr) methods were previously described (5–17). Briefly, time series were demeaned and normalized, followed by generating dense connectomes correlating each vertex/voxel with all other vertices/voxels in the CIFTI grayordinates, and then transformed to Fisher *z* values. For each vertex/voxel, GBCr is calculated as the standardized (*z*-scored) average across those Fisher *z* values with parcel-constrained smoothing (sigma = 4.2 mm), which generates a map for each *f*MRI session where each vertex/voxel value represents the functional connectivity strength of that grayordinate with the rest of the brain. In graph theory terms, GBCr (also known as Functional Connectivity Strength (18)) is considered a weighted measure of nodal strength of a voxel in the whole brain network – determining brain hubs and examining the coherence between a local region and the rest of the brain (19).

**Figure S1.**
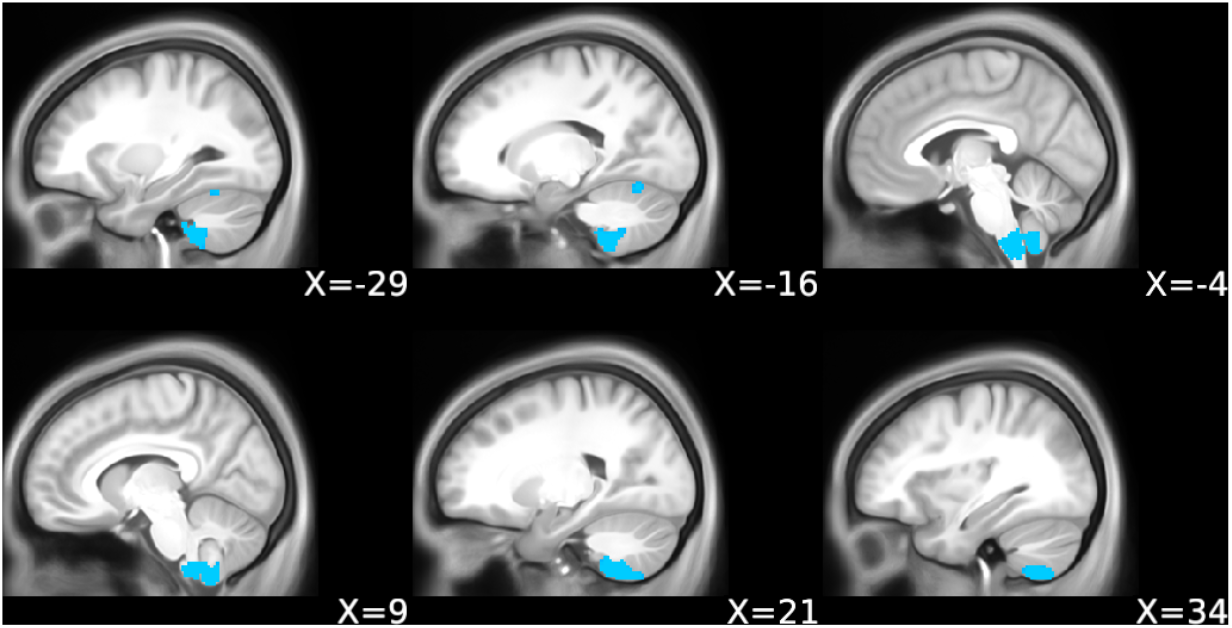
Cerebellar Global Connectivity Post-treatment. The blue clusters mark the vertices with reduced global brain connectivity with global signal regression (GBCr) in posttraumatic stress disorder (PTSD) compared to controls during symptom provocation (*p* < .005 and corrected *α* = .05).

Similar to previous studies (5–12,14,18,20), we have used GBCr, instead of GBC without global signal regression (GBCnr), because the study hypotheses were based on previous GBCr findings (6–8), which provided the rationale for the current report and will facilitate the interpretation of the study findings (see Ref (7) for additional justification). In addition, previous work underscored the need for MGTR to adequately minimize spurious artifacts (4).

### Cerebellar Clusters

Using a GLM comparable to the salience and executive ROIs, we found significant effects of group (*F*_(2,55)_ = 7.2, *p* = .002), time (*F*_(1,55)_ = 25.8, *p* < .001), task (*F*_(1,55)_ = 79.2, *p* < .001), and task*time (*F*_(1,55)_ = 47.3, *p* < .001), task*group (*F*_(2,55)_ = 9.1, *p* < .001), and time*group interactions (*F*_(2,55)_ = 25.8, *p* < .001) on the cerebellar ROI. There were no time*group and no time*task*group interactions (all *p* values > .1). The post-hoc analyses results are shown in Fig. S2.

**Figure S2.**
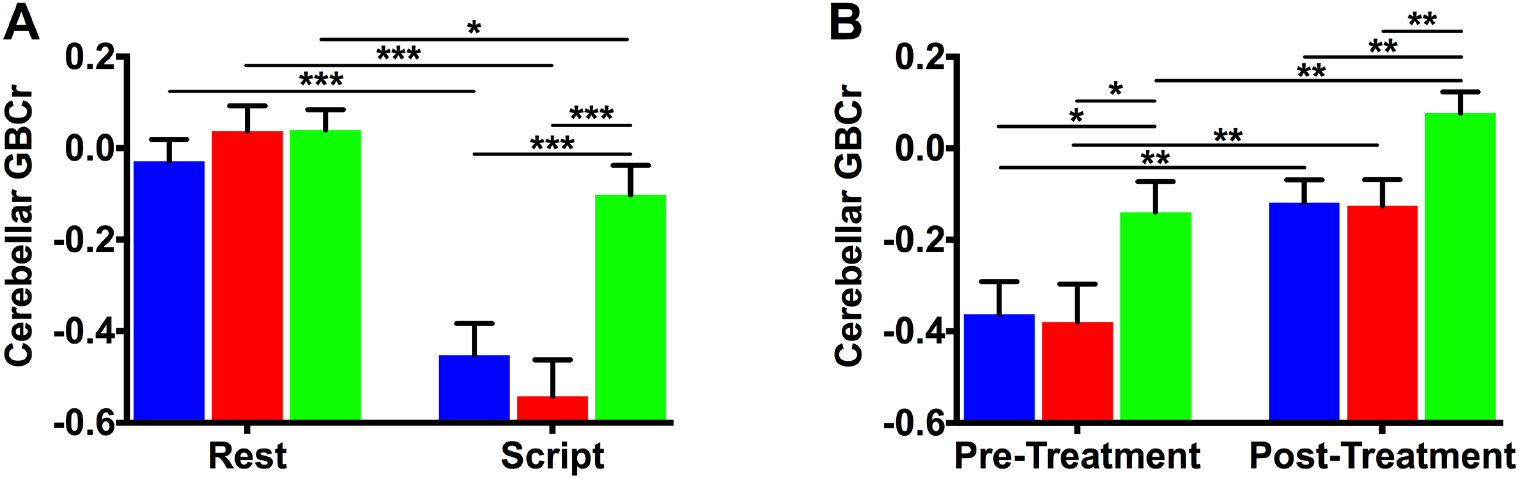
The Effects of Evidence-based Psychotherapy on Cerebellar Connectivity. **A.** There was a significant group by task interaction effect on executive global brain connectivity with global signal regression (GBCr), with more pronounced reduction in GBCr during trauma recollection (i.e., script imagery) compared to during resting state in posttraumatic stress disorder (PTSD) patients treated with present-centered therapy (PCT; blue) or cognitive processing therapy (CPT; red), compared to combat control (CC; green). The lower GBCr values in PTSD compared to CC were significant only during trauma recollection, but not a rest. **B**. There was no time by group interaction, such as the increase in GBCr posttreatment was comparable regardless of group affiliation. * *p* ≤ .05; ** *p* ≤ .01; *** *p* ≤ .001.

